# Pharmacological Inhibition of SUMOylation with TAK-981 Mimics Genetic HypoSUMOylation in Murine Perigonadal White Adipose Tissue

**DOI:** 10.1101/2024.11.23.624974

**Authors:** Damien Dufour, Xu Zhao, Florian Chaleil, Patrizia Maria Christiane Nothnagel, Magnar Bjørås, Anne-Marie Lefrançois-Martinez, Antoine Martinez, Pierre Chymkowitch

## Abstract

Post-translational modification by the small ubiquitin-like modifier (SUMO) is essential for cellular differentiation and homeostasis. Here, we investigate the role of SUMOylation in adipose tissue development using TAK-981, a pharmacological inhibitor of SUMOylation. Administration of TAK-981 to mice resulted in significant defect in weight gain and adipocyte atrophy in perigonadal white adipose tissue (gWAT) depots. Gene expression analyses revealed a marked downregulation of adipogenic genes, including *Pparg, Cebpa*, and *Fasn*. Our data thus indicate that TAK-981 treatment impaired adipogenesis in gWAT, consistent with prior findings that SUMOylation supports transcriptional regulation of adipogenesis and lipid metabolism. We also found significant infiltration of immune cells and efferocytosis in gWAT. Our results thus indicate that SUMOylation inhibition using a small molecule phenocopies genetic hypoSUMOylation models, highlighting its critical role in maintaining adipocyte functionality and immune environment. These findings provide evidence that SUMOylation is essential for fat accumulation *in vivo*. Furthermore, given that TAK-981 is currently under clinical evaluation for the treatment of solid tumors, our results underscore the importance of considering the potential unintended effects of SUMOylation inhibition on adipose tissue in patients.

## Introduction

Post-translational modification by the small ubiquitin-like modifier (SUMO) involves the reversible conjugation of the SUMO peptide to target proteins. Over the past decade, extensive research has explored the role of the SUMOylation pathway in cellular differentiation, utilizing a variety of cellular and in vivo models. While the precise mechanisms regulated by SUMOylation and the specificity of its target proteins remain incompletely understood, evidence strongly indicates that a dynamic SUMOylation/deSUMOylation cycle is critical for the development and functionality of numerous cell types and organs [1-3].

SUMOylation plays a crucial role in maintaining the cellular identity of both stem cells and differentiated cells by preventing undesired cellular identity conversions [4-6]. To uncover the SUMO-regulated mechanisms underlying cellular differentiation and fate determination, efforts were made to characterized SUMO targets and SUMO-chromatin landscapes in various biological systems, including mouse organs, embryonic stem cells (ESCs), mouse embryonic fibroblasts (MEFs), and adipocytes, using mass spectrometry and ChIP-sequencing approaches [2,7,8]. These studies have demonstrated that SUMO targets are predominantly nuclear and that chromatin-bound SUMO associates with proteins involved in chromatin organization, epigenetic regulation, and transcriptional control. Furthermore, the application of specific SUMOylation inhibitors, such as ML-792 and TAK-981, has revealed that SUMOylation can either promote or repress transcription, depending on the cellular context [9,10].

Numerous *in vitro* and *in vivo* studies highlight the critical role of the SUMOylation / deSUMOylation pathway in adipose tissue formation. Mice deficient in *Sumo-1* or *Senp2* exhibit resistance to high-fat diet (HFD)-induced obesity [11,12]. Similarly, the targeted loss of *Ubc9* in white adipose tissue results in adipocyte atrophy [13], while the absence of *Senp7* impairs lipid droplet maturation [14]. Consistently, cellular models with loss of *Sumo-1, Ubc9, Senp1*, or *Senp2* demonstrate impaired adipogenesis, accompanied by disrupted expression of adipogenic genes and genes involved in lipid and energy metabolism [15-17]. These findings underscore the essential role of an active SUMOylation / deSUMOylation cycle in regulating adipose tissue development and metabolic homeostasis.

Given the limited understanding of the mechanisms by which SUMOylation supports adipogenesis, we recently employed adipocyte differentiation as a model to investigate how the SUMOylation pathway regulates differentiation and fat accumulation *in vitro*. Our findings revealed that the SUMOylation pathway is strongly activated upon adipogenic induction and plays a dual role: repressing pre-adipogenic genes while activating adipogenic genes [2]. Mass spectrometry data identified complexes associated with SUMO-dependent heterochromatin formation and DNA methylation, contributing to the repression of preadipocyte-specific genes. These results align with prior studies [7]. Additionally, data from mass spectrometry, ChIP-sequencing, and prolonged treatment with the SUMOylation inhibitor ML-792 demonstrated that SUMOylation enhances the specificity of PPARγ-RXR binding to their target DNA sequences, ensuring the proper expression of target genes critical for adipogenesis. These findings corroborate previously reported roles of SUMOylation in transcriptional regulation [18]. Collectively, our results indicated that an active SUMOylation pathway is indispensable for full adipocyte differentiation, highlighting a unique adipogenesis-specific function of SUMOylation in contrast to other cellular systems [4,5]. Epididymal white adipose tissue (eWAT) undergoes postnatal development, with the first adipocytes observed by postnatal day 7 (p7) [19]. This is followed by simultaneous lipogenesis and adipogenesis, contributing to the tissue’s growth and maturation. The post-weaning period, occurring after p21, represents a critical window for adipose tissue remodeling and is particularly responsive to environmental influences [20,21]. This period presents a unique opportunity to investigate the effects of hypoSUMOylation on eWAT development.

Based on the lipoatrophic effects observed with pharmacological SUMOylation inhibition *in vitro*, we hypothesized that inhibiting SUMOylation using the small-molecule SAE inhibitor TAK-981 would induce adipocyte atrophy not only in eWAT but also across other fat depots, which would underscore the significance of SUMOylation in maintaining adipocyte function and adipose tissue homeostasis during critical developmental stages.

## Results

### SUMOylation Inhibition Induces Adipocyte Atrophy in gWAT

To evaluate the impact of pharmacological inhibition of SUMOylation on adipose tissue *in vivo*, we administered the SUMOylation inhibitor TAK-981 [22,23] to mice, following a previously established protocol [1]. TAK-981 was administered biweekly over a period of five weeks. Body mass was monitored throughout this period, revealing that, consistently with a previous cohort [1], male mice treated with TAK-981 exhibited a significantly attenuated rate of weight gain compared to untreated controls, with the statistical significance of this difference increasing over time (Figure 1a).

**Figure 1.**
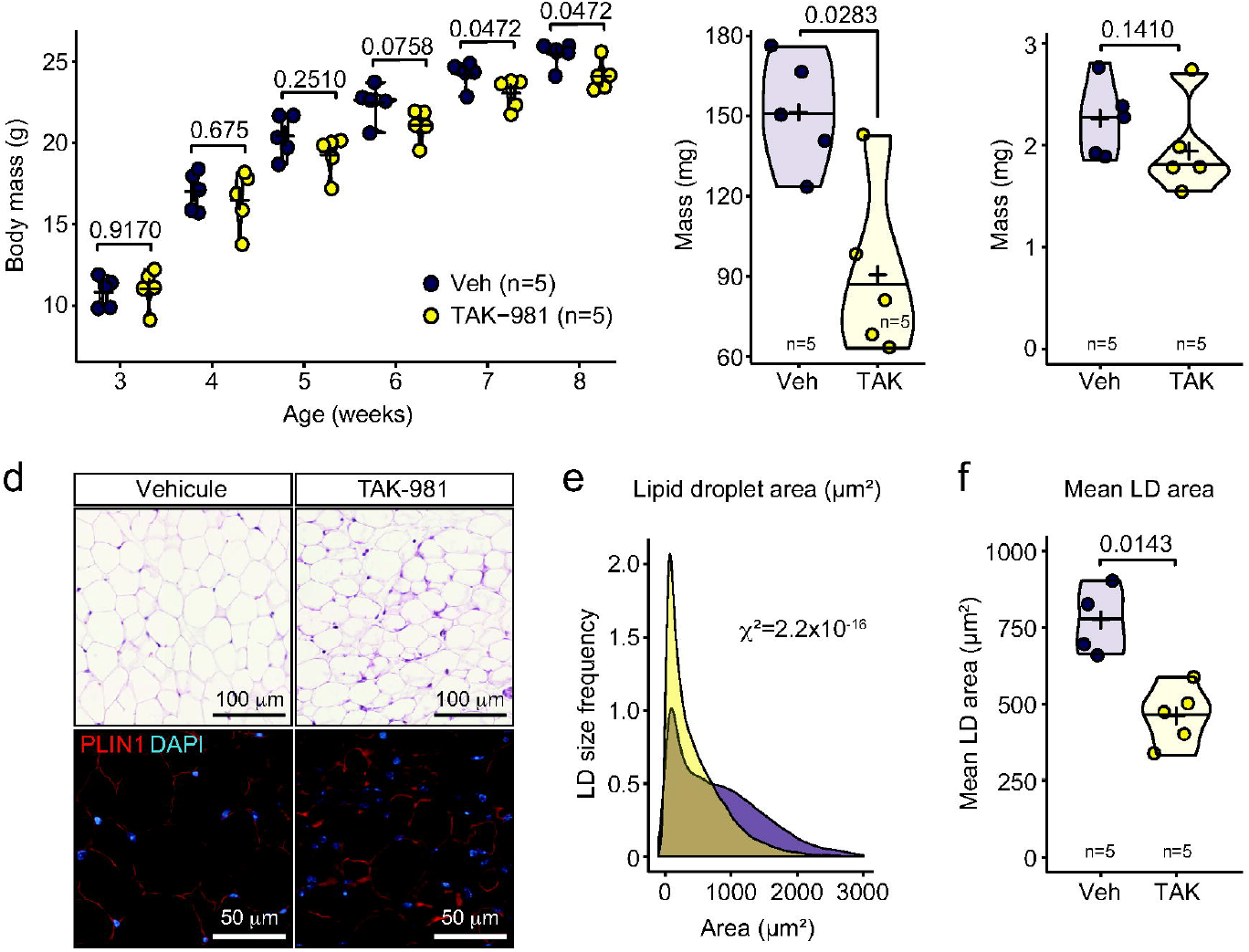
SUMOylation inhibition induces eWAT atrophy. ***Let me know if you’d like this adjusted further!*** (a) body mass follow-up of male mice treated with vehicle or TAK-981 between 3 and 8 weeks of age. (b-c) absolute mass of eWAT *(b)* and adrenal glands *(c)* at 8 weeks of age. (d-f) H&E staining (top) and PLIN1 immunofluorescence (bottom) *(d)* and quantification of lipid droplet area distribution *(e)* and lipid droplet mean area *(f)* of eWAT of 8-week-old mice. **Figure 1 alt text: Impact of SUMOylation inhibition on eWAT development**. (a) Line graph showing body mass follow-up in male mice treated with vehicle or TAK-981 from 3 to 8 weeks of age. (b-c) Bar graphs presenting the absolute mass of eWAT (b) and adrenal glands (c) at 8 weeks of age. (d) Images of eWAT tissue from 8-week-old mice, with H&E staining (top) and PLIN1 immunofluorescence (bottom). (e) Graph showing lipid droplet area distribution in eWAT. (f) Bar graph depicting lipid droplet mean area in eWAT. TAK-981 treatment is associated with eWAT atrophy and changes in lipid droplet morphology.”

The reduced weight gain observed in TAK-981-treated mice was associated with adipocyte atrophy, as evidenced by a decrease in perigonadal white adipose tissue (gWAT) mass, specifically epididymal WAT (eWAT), while adrenal gland mass remained unaffected (Figure 1b and 1c). To further investigate the cause of eWAT hypoplasia, we analyzed the size of lipid droplets within the tissue. Lipid droplets in TAK-981-treated mice were significantly smaller than those in control mice (Figure 1d-f).

We subsequently investigated whether adipocyte atrophy was restricted to eWAT or occurred across other fat depots in female mice. Although one mouse of the cohort did not respond to the treatment, similar adipocyte atrophy was observed in periovarian white adipose tissue (poWAT) (Figure 2a and b). However, inguinal white adipose tissue (iWAT) and interscapular brown adipose tissue (iBAT) remained unaffected (Figure 2a and b). These results indicate that pharmacological inhibition of SUMOylation by TAK-981 induces adipocyte atrophy and reduces white adipose tissue mass in gWAT.

**Figure 2.**
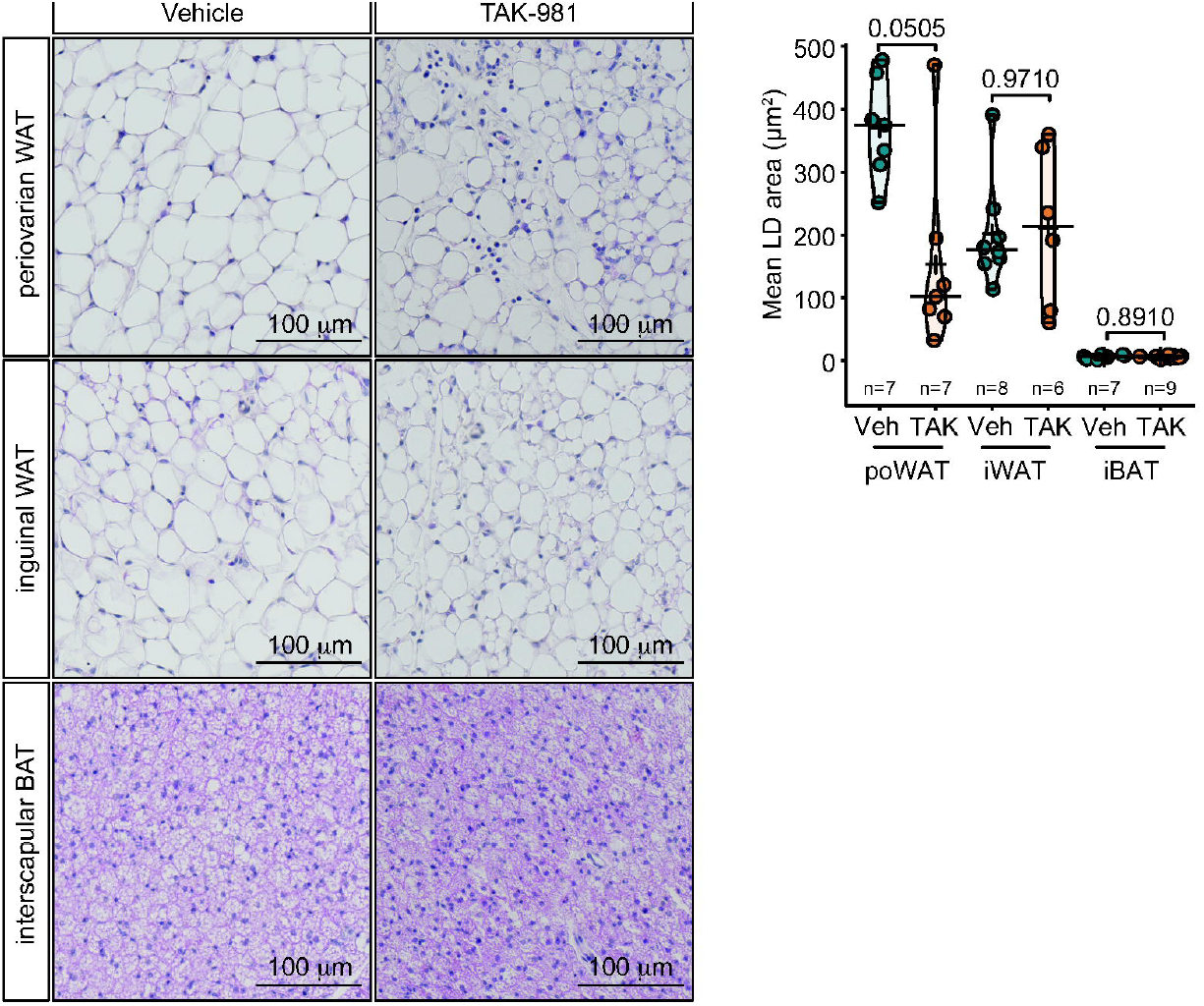
TAK-981 affects poWAT in female mice. (a) H&E staining of periovarian WAT (poWAT), inguinal WAT (iWAT) and interscapular BAT (iBAT) in female mice treated with vehicle or TAK-981. (b) Quantification of mean lipid droplet area in poWAT, iWAT and iBAT. **Figure 2 alt text: Impact of TAK-981 on different WAT depots in female mice**. (a) Representative images of H&E-stained periovarian WAT (poWAT), inguinal WAT (iWAT), and interscapular BAT (iBAT) from female mice treated with vehicle or TAK-981. (b) Bar graphs quantifying the mean lipid droplet area in poWAT, iWAT, and iBAT. TAK-981 treatment influences lipid droplet morphology in these tissues.

### SUMOylation inhibition leads to downregulation of adipogenic genes

The results presented above suggested two non-exclusive hypotheses explaining adipocyte atrophy in gWAT.

First, gWAT could undergo beiging. However, we found no multilocular adipocytes in TAK-981-treated mice, and we were unable to detect UCP1-positive cells in eWAT under either treatment condition (Figure 3a). Additionally, the expression of *Ucp1* and *Cidea*, two beige adipocyte markers, was unaffected by TAK-981 treatment, as demonstrated by qPCR analysis (Figure 3b). These data indicated that inhibiting SUMOylation did not induce beiging in eWAT.

**Figure 3.**
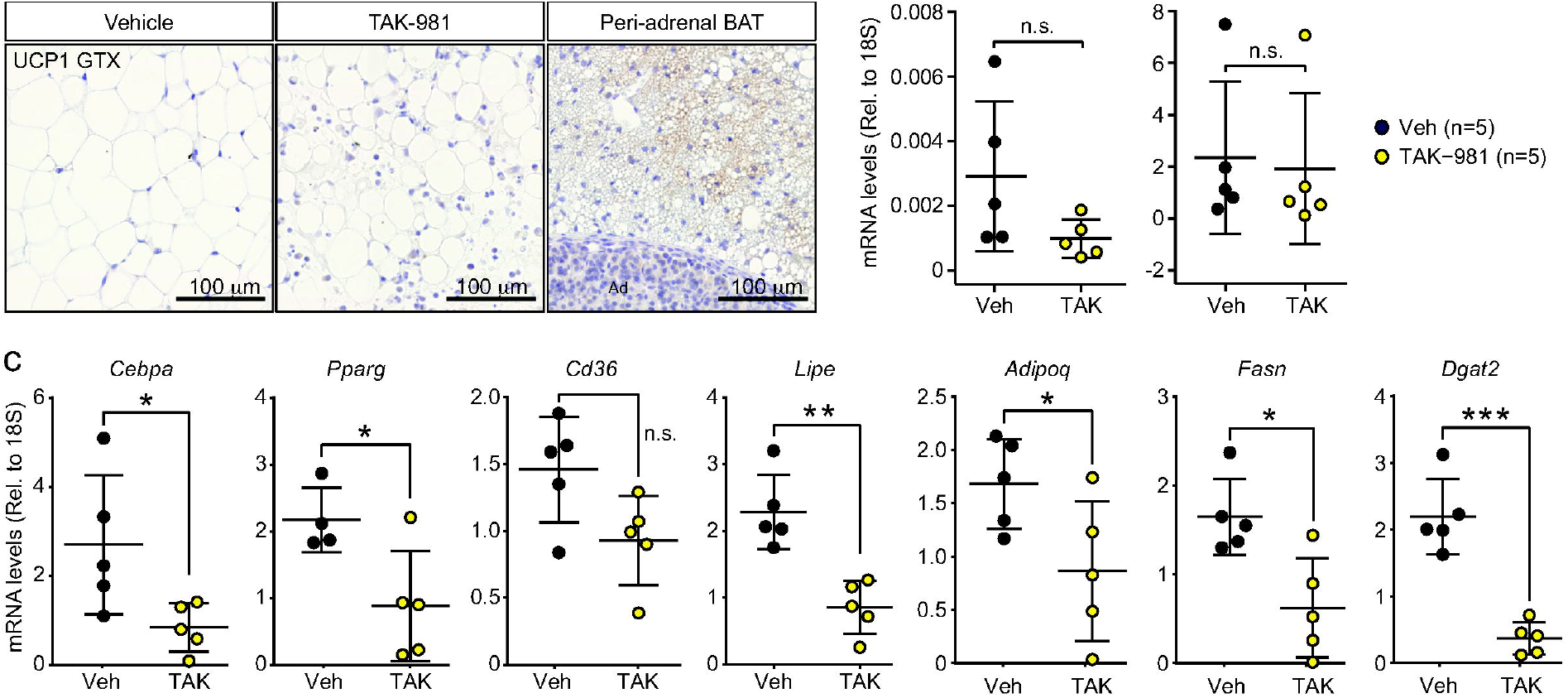
Downregulation of adipogenic genes in TAK-981-treated mice. (a) immunofluorescence staining of UCP1 in eWAT from vehicle or TAK-981-treated mice. (b) RT-qPCR analysis of genes associated with beige identity *Cidea* and *Ucp1*. (c) RT-qPCR analysis of genes involved in adipogenesis *Cebpa, Pparg, Cd36, Lipe, Adipoq, Fasn and Dgat2*. *: *P* values ≤ 0.05, **: *P* values ≤ 0.01, ***: *P* values ≤ 0.001. n.s.: not significant. **Figure 3 alt text: Downregulation of adipogenic genes in TAK-981-treated mice**. (a) Immunofluorescence images showing UCP1 expression in eWAT from vehicle- and TAK-981-treated mice. (b) Bar graph displaying RT-qPCR analysis of beige identity-related genes, Cidea and Ucp1. (c) Bar graph showing RT-qPCR analysis of adipogenesis-associated genes, including Cebpa, Pparg, Cd36, Lipe, Adipoq, Fasn, and Dgat2. Statistical significance indicated by P values (* ≤ 0.05, ** ≤ 0.01, *** ≤ 0.001, n.s.: not significant).

Second, as demonstrated in our previous studies with mouse 3T3-L1 cells, prolonged SUMOylation inhibition correlates with the downregulation of adipogenic genes, indicating a potential defect in adipogenesis [2]. To explore this in mice, we purified RNA from eWAT collected from mice treated with or without TAK-981 and assessed the expression of adipogenic genes using RT-qPCR. This analysis revealed that the expression of key WAT markers, namely *Cebpa, Pparg, Cd36, Lipe, Adipoq, Fasn*, and *Dgat2*, was significantly downregulated in TAK-981-treated mice (Figure 3c). These findings indicate that TAK-981 inhibits adipogenic transcription *in vivo*.

This result suggests that adipocyte differentiation is impaired by TAK-981, aligning with our observations in 3T3-L1 cells and corroborating the results presented in Figure 1 [2].

Together, these data reinforce the hypothesis that prolonged SUMOylation inhibition disrupts the transcriptional program necessary for adipogenesis.

### Immune cells infiltration in gWAT

Previous studies indicated that weight loss, adipocyte atrophy and adipogenic defects induce immune cells invasion in WAT [24]. Furthermore, accumulation of immune cells has been observed upon genetic HypoSUMOylation in adipocyte-specific *Ubc9* cKO mice [13]. Further examination of eWAT and poWAT morphology revealed the accumulation of small cells with round nuclei located between adipocytes resembling immune cells (Figure 4). To test whether SUMOylation inhibition triggered accumulation of immune cells, we assessed the presence of leukocyte marker CD45 and macrophage marker IBA1 in eWAT using immunohistofluorescence. Quantification revealed an increase in the proportion of both CD45+ cells and IBA1+ cells in male mice treated with TAK-981 (Figure 4a-c). Consistently, we detected the accumulation of IBA1+ cells in poWAT in female mice (Figure 4d). This accumulation was confirmed by increased mRNA levels of *Cd68*, a macrophage marker, in the eWAT of mice treated with TAK-981 when compared to untreated animals (Figure 4e). Finally, we observed a significant increase of the presence of IBA1+ crown-like figures indicating active efferocytosis in eWAT (Figure 4a and f), a process where macrophages remove stressed or damaged adipocytes through phagocytosis [25]. To assess this, we used γH2A.X staining as a marker of double strand DNA damage repair, hallmark of damaged, apoptotic or senescent cells [26,27]. Our data indicated an increase in the proportion of nuclei containing γH2A.X foci in TAK-981-treated mice (Figure 5a and b). Together these results show that low SUMOylation caused by TAK-981 treatment triggers an alteration of adipocytes physiology associated with a macrophage infiltration probably involved in efferocytosis in mouse gWAT.

**Figure 4.**
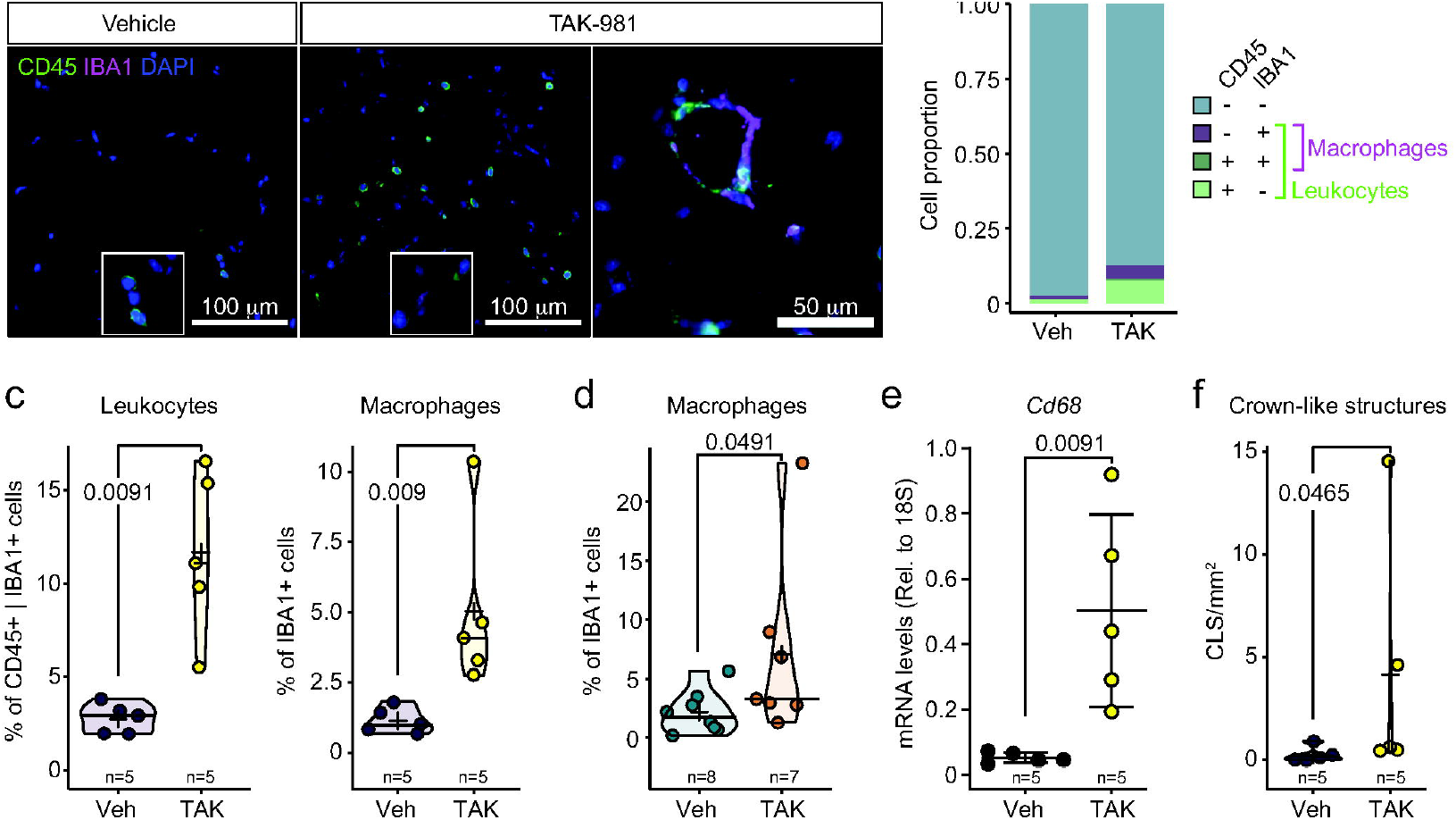
Immune cells infiltration in gWAT upon SUMOylation inhibition. (a) Coimmunofluorescent labelling of pan leukocyte marker CD45 (green) and macrophage marker IBA1 (purple) in 8-week-old eWAT from mice treated with vehicle or TAK-981. (b) Repartition of macrophages and other leukocytes amongst eWAT cells. (c) Quantification of macrophages (IBA1+) and total leukocytes (CD45+ and/or IBA1+) in eWAT. (d) Quantification of macrophages (IBA1+) in poWAT. (e) RT-qPCR analysis of the macrophage marker *Cd68* in eWAT from mice treated with vehicle or TAK-981. (f) Quantification of crown-like structures density in eWAT. **Figure 4 alt text: Immune cell infiltration in gWAT following SUMOylation inhibition**. (a) Coimmunofluorescence images showing CD45 (pan-leukocyte marker, green) and IBA1 (macrophage marker, purple) in eWAT from 8-week-old mice treated with vehicle or TAK-981. (b) Pie chart depicting the distribution of macrophages and other leukocytes in eWAT. (c) Bar graphs quantifying macrophages (IBA1+) and total leukocytes (CD45+ and/or IBA1+) in eWAT. (d) Bar graph quantifying macrophages (IBA1+) in poWAT. (e) Bar graph showing RT-qPCR analysis of the macrophage marker Cd68 in eWAT. (f) Bar graph illustrating the density of crown-like structures in eWAT. TAK-981 treatment is associated with increased immune cell infiltration and macrophage activity in gWAT

**Figure 5:**
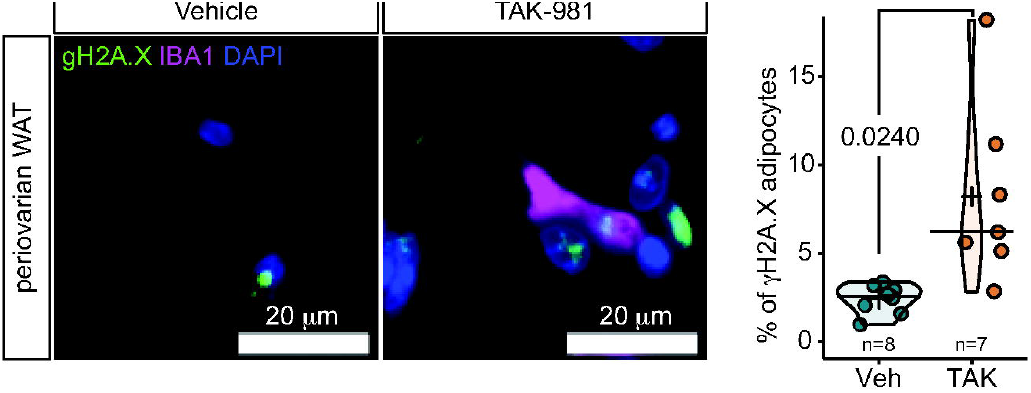
Immune cells infiltration is associated with DNA damage in adipocytes. (a) Representative picture of coimmunofluorescent labelling of γH2A.X (green) and IBA1 (purple) in 8-week-old poWAT. (b) Quantification of IBA1 negative cells harbouring γH2A.X foci in poWAT from mice treated with vehicle or TAK-981. **Figure 5 alt text: Immune cell infiltration associated with DNA damage in adipocytes. (a)** Representative coimmunofluorescence image showing γH2A.X (DNA damage marker, green) and IBA1 (macrophage marker, purple) in poWAT from 8-week-old mice. (b) Bar graph quantifying IBA1-negative cells with γH2A.X foci in poWAT from vehicle- and TAK-981-treated mice. TAK-981 treatment correlates with increased DNA damage in non-macrophage cells.

## Discussion

Recent studies using various cellular and *in vivo* models have consistently demonstrated that SUMOylation plays a crucial role in limiting cellular differentiation, favoring stemness, and maintaining cellular identity over reprogramming [1,4,5]. However, other research has indicated that the formation and function of adipose tissue require an active SUMOylation pathway. For instance, *Sumo-1* KO mice and conditional KO of *Ubc9* in WAT exhibit resistance to high-fat diet-induced obesity and WAT atrophy, respectively [11,13,28]. Moreover, studies from our lab and others have shown that maintaining low SUMOylation levels in differentiating mouse 3T3-L1 cells disrupts adipogenic gene expression dynamics, leading to adipocyte atrophy [2,15]. These findings suggest that adipocyte differentiation is specifically impaired by reduced SUMOylation.

To explore this further, we treated mice with TAK-981, a potent inhibitor of SUMOylation. Pharmacological inhibition of SUMOylation with TAK-981 resulted in significant atrophy of eWAT and poWAT, a consequence primarily attributed to adipocyte atrophy [29]. This further supports the notion that an active SUMOylation pathway is essential for proper adipocyte differentiation and adipose tissue homeostasis.

The anti-differentiation effect of SUMOylation has primarily been observed at the stem cell level, where high SUMOylation levels inhibit differentiation [1,4,5]. Since adipose tissue contains approximately 5% mesenchymal adipose stem cells, it is possible that TAK-981, while reducing overall eWAT mass, may exert a pro-adipogenic effect on these stem cells, potentially leading to an increase in the number of adipocytes. This possibility is plausible, as eWAT undergoes both lipogenesis and adipogenesis between p7 and adulthood [30], and will be investigated in a subsequent study.

However, our data suggest that, at the whole tissue level, pharmacological inhibition of SUMOylation primarily affects mature adipocytes, which constitute most cells in WAT. We propose that TAK-981-induced adipocyte atrophy results from the inability of differentiated adipocytes to accumulate fat. While the effect on stem cells cannot be entirely ruled out, it likely remains undetected due to the predominance of mature adipocytes in the tissue. This is further supported by the downregulation of adipogenic genes (e.g., *Pparg, Fasn*), suggesting a failure in adipogenesis, likely caused by alterations in chromatin structure, transcriptional activity, and epigenetic regulation. These observations are consistent with our previous findings in 3T3-L1 adipocytes [2].

In this context, SUMOylation inhibition likely destabilizes chromatin-bound complexes and negatively impacts transcriptional regulators, as has been described in previous studies [2,31-33]. Our data underscore the critical role of SUMOylation in maintaining proper adipogenic gene expression and adipocyte functionality.

Furthermore, previous studies have directly linked SUMOylation to the maintenance of metabolic pathways involved in fatty acid synthesis [34]. These studies suggest that, in addition to regulating the expression of key genes such as *Fasn*, inhibition of SUMOylation could also directly impact the activity of enzymes responsible for fatty acid synthesis. Our recent mass spectrometry analysis in differentiating 3T3-L1 cells further supports this notion, revealing that enzymes involved in fatty acid synthesis, such as FASN, are SUMO targets [2]. This suggests that SUMOylation not only regulates the transcription of adipogenic genes but may also modulate the enzymatic activity of proteins critical for lipid metabolism. Therefore, SUMOylation likely plays a central role in coordinating both the expression and activity of enzymes involved in fatty acid synthesis, thereby contributing to adipocyte differentiation and lipid accumulation.

Although this study focused on adipose tissue, this organ is particularly sensitive to the energy state of the organism. As observed in adipocyte-specific *Ubc9* KO, it is very likely that energy balance is affected following TAK-981 treatment. However, the consistency with cellular models and the similarity of our model with the adipose tissue-specific ablation of *Ubc9* [13] support the adipose origin of the phenotype. Similarly, even though we did not observe TAK-981 induced beiging, adipocytes from male eWAT do not seem to be able to transdifferentiate into a beige identity [35] and it cannot be excluded that thermogenesis took place in the iWAT or BAT after TAK-981 treatment even though there was no apparent change in iWAT or BAT morphology. Moreover, the absence of beiging in TAK-981 treated mice is consistent with the observation that beiging of iWAT is dependent on active SUMOylation as evidenced by adipose tissue-specific *Senp2* KO [36].

These *in vivo* data suggest that the administration of TAK-981, which is currently in phase 1-2 clinical trials for solid tumors [22,37], to individuals could result in weight loss and therefore constitute an inroad into the development of new anti-obesity drugs. However, despite the apparent negative effect of TAK-981 on fat accumulation, this strategy is not viable at this stage. First, adipocyte atrophy with constant energy intake is not sought to fight obesity because fatty acids will in return accumulate in other organs such as the liver and lead to metabolism-associated fatty liver disease [38]. Second, low SUMOylation levels in WAT correlated with the infiltration of lymphocytes and activated macrophages, and efferocytosis, which suggests activation of the immune system as previously shown [13,39]. However, considering that TAK-981 is currently being evaluated in clinical trials for solid tumors, our data provide valuable insights into the potential unintended effects of this compound in patients.

Overall, we show that the pharmacological reduction of global SUMOylation phenocopies both conditional [13] and total [20] genetic hypoSUMOylation models, making it an interesting tool to study generalized adipocyte atrophy syndroms. TAK-981 allows easy manipulation of the SUMOylation machinery during various stage of post-natal development and aging. However, authors wanting to use TAK-981 to study the result of hypoSUMOylation should consider the potential effects of increased circulating free fatty acids resulting from adipocyte atrophy in metabolic organs [40].

## Experimental procedures

### Ethical approval declarations

Mouse experiments were conducted according to French and European directives for the use and care of animals for research purposes and were approved by the Comité d’Éthique pour l’Expérimentation Animale en Auvergne (project agreement #211522019061912052883), C2EA-02, at Institut National de Recherche pour l’Agriculture, l’Alimentation et l’Environnement, Research Centre Clermont-Theix, France (C2E2A). We have adhered to ARRIVE guidelines and upload a completed checklist.

### Animals

Mice were obtained from Janvier Labs (France) and then bred in-house and maintained on a mixed sv129-C57Bl/6 genetic background were housed on a 12-h light/12-h dark cycle (lights on at 7:00 am). Mice were fed normal, commercial rodent chow and provided with water ad libitum. After weaning, mice were kept in siblings with a maximum of four animals per cage. At the end of experimental procedures mice were killed by decapitation around 8:30 am and trunk blood was collected in vacuum blood collection tubes (VF-053STK, Terumo). Mice aged 3 weeks were treated with vehicle control or TAK-981 (MedChemExpress, Sollentuna, Sweden) (7.5 mg/kg, intraperitoneally twice per week) for 5 weeks as previously described [1]. According to the ethical approval mentioned above, at the end of experimental procedures mice were killed by decapitation to prevent stress.

### Histology

Tissues were fixed in 4% PFA for 24h and embedded in paraffin. 5 μm sections were deparaffinized and processed for hematoxylin/eosin staining. For immunofluorescence, deparaffinized slides were submerged in a citrate-based antigen retrieval buffer and microwaved for 8 min. After being rinsed with 1x PBS, slides were blocked for an hour with 2.5% horse serum (Vector) and incubated overnight at 4C with rabbit anti-PLIN1 antibody (Cell signaling technology #9349, 1/500), anti-rat CD45 (550539 1/200), anti-mouse γH2A.X (Merck 05-636) or anti rabbit IBA1 (ABIN2857032 1/500). After rinsing, they were incubated with ImmPRESS polymer for 30 min at room temperature. HRP activity was detected with Alexafluor (Thermo Fisher). Images were acquired with Zeiss Axioscan Z1 and ZEN 3.4 blue edition software and analyzed with QuPath 0.4.3 software [41]. Lipid droplet quantification was performed on PLIN1 staining (Figure 1) or H&E staining (Figure 2) using QuPath 0.4.3 software (https://github.com/Damien-Dufour/Image_analysis).

### RNA purification, cDNA synthesis and RTqPCR

Total RNA purification from Formalin-Fixed Paraffin-Embedded eWAT was done using the RNeasy FFPE kit (73504, Qiagen) and cDNA synthesis was done using the QuantiTect Reverse Transcription Kit (205311, Qiagen) following manufacturer’s instructions. RT-qPCR was performed using the HOT FIREPol® EvaGreen® qPCR Supermix (Solis Biodyne, 08-36-00001-5) on a LightCycler 96 instrument (Roche Diagnostics). Gene expression was normalized to the expression of *18S* and plots and data analyzed and visualized using GraphPad Prism 9. *P-*values are indicated in figure legends. The following primer pairs were used for RT-qPCRs.

*Cebpa*: CAAAGCCAAGAAGTCGGTGGACAA – TCATTGTGACTGGTCAACTCCAGC

*Pparg*: CCGTAGAAGCCGTGCAAGAG – CCGTAGAAGCCGTGCAAGAG

*Cd36*: CGACTGCAGGTCAACATATT – TTAGCCACAGTATAGGTACA

*Lipe*: GCTCATCTCCTATGACCTACGG – TCCGTGGATGTGAACAACCAGG

*AdipoQ*: CTTGTGCAGGTTGGATGGCA – GGACCAAGAAGACCTGCATCT

*Fasn*: CACAGTGCTCAAAGGACATGCC – CACCAGGTGTAGTGCCTTCCTC

*Dgat2*: CTGTGCTCTACTTCACCTGGCT – CTGGATGGGAAAGTAGTCTCGG

*Cidea*: TGCTCTTCTGTATCGCCCAGT – GCCGTGTTAAGGAATCTGCTG

*Ucp1*: ACTGCCACACCTCCAGTCATT – CTTTGCCTCACTCAGGATTGG

*Cd68*: CACAGTGGACATTCATGGCG – TGGTCACGGTTGCAAGAGAA

*18S*: GTAACCCGTTGAACCCCATT – CCATCCAATCGGTAGTAGCG

### Statistics

Statistics were conducted using R language (4.2.2) and Comp3Moy function from sumo package (https://github.com/Damien-Dufour/sumo) or GraphPad Prism (v7). Normality of populations distribution was assessed with Shapiro & Wilk test for n > 7 or Kolmogorov– Smirnov normality test. If data followed a normal distribution, homoscedasticity was estimated with a Barlett test. To compare two populations, unpaired, two-tailed t test was used for normally distributed data with the same variance, Mann–Whitney for non-normal distributions and Welch t test for normally distributed data but with different variances. To compare three or more distributions: one-way ANOVA for normally distributed samples with pairwise multiple t tests or Kruskal–Wallis for nonnormally distributed samples with planned comparisons using Dunn’s test to determine the genotype effect or the treatment effect. Crosses on the violin plots represent the mean and lines represent the median. Error bars in scatterplots represent the SD. Number of samples is indicated at the bottom of the graph.

## Author contributions

**Damien Dufour:** Methodology, Investigation, Formal analysis, Visualization, Writing - Original Draft. **Xu Zhao:** Investigation, Formal analysis, Visualization. **Florian Chaleil:** Investigation. **Patrizia Maria Christiane Nothnagel:** Investigation, Formal analysis, Visualization. **Magnar Bjørås:** Funding acquisition, Resources. **Anne-Marie Lefrançois-Martinez:** Supervision, Investigation. **Antoine Martinez:** Supervision, Funding acquisition, Resources, Writing - Review & Editing. **Pierre Chymkowitch:** Conceptualization, Resources, Supervision, Funding acquisition, Writing - Original Draft, Writing - Review & Editing. All authors have read and approved the final work.

## Funding and additional information

This research was supported by the Research Council of Norway (grant numbers 353961 to P.C. and 326101 to M.B.), with additional institutional support provided by the Department of Biosciences, University of Oslo and UNIFOR (to P.C.). Further institutional support was received from CNRS, INSERM, Université Clermont-Auvergne, and grants from the Ministère de l’Enseignement Supérieur, de la Recherche et de l’Innovation (to D.D.), the Société Française d’Endocrinologie (to D.D. and A.M.), and the Agence Nationale de la Recherche (grant number ANR-18-CE14-0012-02 to A.M.).

## Data accessibility

Data can be accessed at figshare.com https://doi.org/10.6084/m9.figshare.28008926.

## References

1. Dufour D, Dumontet T, Sahut-Barnola I, et al. Loss of SUMO-specific protease 2 causes isolated glucocorticoid deficiency by blocking adrenal cortex zonal transdifferentiation in mice. Nat Commun. 2022 Dec 21;13(1):7858.

2. Zhao X, Hendriks IA, Le Gras S, et al. Waves of sumoylation support transcription dynamics during adipocyte differentiation. Nucleic Acids Res. 2022 Feb 22;50(3):1351–1369.

3. Demarque MD, Nacerddine K, Neyret-Kahn H, et al. Sumoylation by Ubc9 regulates the stem cell compartment and structure and function of the intestinal epithelium in mice. Gastroenterology. 2011 Jan;140(1):286–96.

4. Cossec JC, Theurillat I, Chica C, et al. SUMO Safeguards Somatic and Pluripotent Cell Identities by Enforcing Distinct Chromatin States. Cell Stem Cell. 2018 Nov 1;23(5):742–757 e8.

5. Cossec JC, Traboulsi T, Sart S, et al. Transient suppression of SUMOylation in embryonic stem cells generates embryo-like structures. Cell reports. 2023 Apr 15;42(4):112380.

6. Bjune J-I, Laber S, Lawrence-Archer L, et al. Mechanisms of the <em>FTO</em> locus association with obesity: Irx3 controls a sumoylation-dependent switch between adipogenesis and osteogenesis. bioRxiv. 2023:2023.10.17.562662.

7. Theurillat I, Hendriks IA, Cossec JC, et al. Extensive SUMO Modification of Repressive Chromatin Factors Distinguishes Pluripotent from Somatic Cells. Cell reports. 2020 Sep 15;32(11):108146.

8. Hendriks IA, Lyon D, Su D, et al. Site-specific characterization of endogenous SUMOylation across species and organs. Nat Commun. 2018 Jun 25;9(1):2456.

9. Boulanger M, Chakraborty M, Tempe D, et al. SUMO and Transcriptional Regulation: The Lessons of Large-Scale Proteomic, Modifomic and Genomic Studies. Molecules. 2021 Feb 5;26(4).

10. Chymkowitch P, Nguea PA, Enserink JM. SUMO-regulated transcription: challenging the dogma. Bioessays. 2015 Oct;37(10):1095–105.

11. Mikkonen L, Hirvonen J, Janne OA. SUMO-1 regulates body weight and adipogenesis via PPARgamma in male and female mice. Endocrinology. 2013 Feb;154(2):698–708.

12. Zheng Q, Cao Y, Chen Y, et al. Senp2 regulates adipose lipid storage by de-SUMOylation of Setdb1. J Mol Cell Biol. 2018 Jun 1;10(3):258–266.

13. Cox AR, Chernis N, Kim KH, et al. Ube2i deletion in adipocytes causes lipoatrophy in mice. Mol Metab. 2021 Jun;48:101221.

14. Pei J, Zou D, Li L, et al. Senp7 deficiency impairs lipid droplets maturation in white adipose tissues via Plin4 deSUMOylation. Journal of Biological Chemistry. 2024 2024/06/01/;300(6):107319.

15. Cignarelli A, Melchiorre M, Peschechera A, et al. Role of UBC9 in the regulation of the adipogenic program in 3T3-L1 adipocytes. Endocrinology. 2010 Nov;151(11):5255–66.

16. Liu B, Wang T, Mei W, et al. Small ubiquitin-like modifier (SUMO) protein-specific protease 1 de-SUMOylates Sharp-1 protein and controls adipocyte differentiation. J Biol Chem. 2014 Aug 8;289(32):22358–64.

17. Chung SS, Ahn BY, Kim M, et al. Control of adipogenesis by the SUMO-specific protease SENP2. Mol Cell Biol. 2010 May;30(9):2135–46.

18. Sri Theivakadadcham VS, Bergey BG, Rosonina E. Sumoylation of DNA-bound transcription factor Sko1 prevents its association with nontarget promoters. PLoS Genet. 2019 Feb;15(2):e1007991.

19. Han J, Lee JE, Jin J, et al. The spatiotemporal development of adipose tissue. Development. 2011 Nov;138(22):5027–37.

20. Bruder J, Fromme T. Global Adipose Tissue Remodeling During the First Month of Postnatal Life in Mice. Front Endocrinol (Lausanne). 2022;13:849877.

21. Holtrup B, Church CD, Berry R, et al. Puberty is an important developmental period for the establishment of adipose tissue mass and metabolic homeostasis. Adipocyte. 2017 Jul 3;6(3):224–233.

22. Langston SP, Grossman S, England D, et al. Discovery of TAK-981, a First-in-Class Inhibitor of SUMO-Activating Enzyme for the Treatment of Cancer. Journal of Medicinal Chemistry. 2021 2021/03/11;64(5):2501–2520.

23. Du L, Liu W, Aldana-Masangkay G, et al. SUMOylation inhibition enhances dexamethasone sensitivity in multiple myeloma. Journal of Experimental & Clinical Cancer Research. 2022 2022/01/04;41(1):8.

24. Kosteli A, Sugaru E, Haemmerle G, et al. Weight loss and lipolysis promote a dynamic immune response in murine adipose tissue. J Clin Invest. 2010 Oct;120(10):3466–79.

25. Boutens L, Stienstra R. Adipose tissue macrophages: going off track during obesity. Diabetologia. 2016 May;59(5):879–94.

26. Rogakou EP, Nieves-Neira W, Boon C, et al. Initiation of DNA Fragmentation during Apoptosis Induces Phosphorylation of H2AX Histone at Serine 139 *. Journal of Biological Chemistry. 2000;275(13):9390–9395.

27. Ishaq A, Dufour D, Cameron K, et al. Metabolic memory of dietary restriction ameliorates DNA damage and adipocyte size in mouse visceral adipose tissue. Exp Gerontol. 2018 Nov;113:228–236.

28. Zhang FP, Mikkonen L, Toppari J, et al. Sumo-1 function is dispensable in normal mouse development. Mol Cell Biol. 2008 Sep;28(17):5381–90.

29. Dufour D, Zhao X, Bjoras M, et al. Pharmacological inhibition of SUMOylation recapitulates genetic hypoSUMOylation in the murine epididymal white adipose tissue. bioRxiv. 2024:2024.11.23.624974.

30. Wang QA, Tao C, Gupta RK, et al. Tracking adipogenesis during white adipose tissue development, expansion and regeneration. Nat Med. 2013 Oct;19(10):1338–44.

31. Launonen K-M, Varis V, Aaltonen N, et al. Central role of SUMOylation in the regulation of chromatin interactions and transcriptional outputs of the androgen receptor in prostate cancer cells. Nucleic Acids Research. 2024.

32. Chymkowitch P, Nguea PA, Aanes H, et al. TORC1-dependent sumoylation of Rpc82 promotes RNA polymerase III assembly and activity. Proc Natl Acad Sci U S A. 2017 Jan 31;114(5):1039–1044.

33. Rosonina E, Akhter A, Dou Y, et al. Regulation of transcription factors by sumoylation. Transcription. 2017 Aug 8;8(4):220–231.

34. Sapir A. Not So Slim Anymore-Evidence for the Role of SUMO in the Regulation of Lipid Metabolism. Biomolecules. 2020 Aug 6;10(8).

35. Kim SN, Jung YS, Kwon HJ, et al. Sex differences in sympathetic innervation and browning of white adipose tissue of mice. Biol Sex Differ. 2016;7:67.

36. Lee JS, Chae S, Nan J, et al. SENP2 suppresses browning of white adipose tissues by de-conjugating SUMO from C/EBPbeta. Cell reports. 2022 Feb 22;38(8):110408.

37. Dudek AZ, Juric D, Dowlati A, et al. Phase 1/2 study of the novel SUMOylation inhibitor TAK-981 in adult patients (pts) with advanced or metastatic solid tumors or relapsed/refractory (RR) hematologic malignancies. Journal of Clinical Oncology. 2021 May 20;39(15).

38. Valenti L, Pelusi S. Redefining fatty liver disease classification in 2020. Liver Int. 2020 May;40(5):1016–1017.

39. Lightcap ES, Yu P, Grossman S, et al. A small-molecule SUMOylation inhibitor activates antitumor immune responses and potentiates immune therapies in preclinical models. Sci Transl Med. 2021 Sep 15;13(611):eaba7791.

40. Sorkina E, Chichkova V. Generalized lipoatrophy syndromes. Presse Med. 2021 Nov;50(3):104075.

41. Bankhead P, Loughrey MB, Fernandez JA, et al. QuPath: Open source software for digital pathology image analysis. Sci Rep. 2017 Dec 4;7(1):16878.

